# Optimum Threshold Minimizes Noise in Timing of Intracellular Events

**DOI:** 10.1101/2020.02.14.949891

**Authors:** Sherin Kannoly, Tianhui Gao, Supravat Dey, Ing-Nang Wang, Abhyudai Singh, John J. Dennehy

## Abstract

How the noisy expression of regulatory proteins affects timing of intracellular events is an intriguing fundamental problem that influences diverse cellular processes. Here we use the bacteriophage λ to study event timing in individual cells where cell lysis is the result of expression and accumulation of a single protein (holin) in the *Escherchia coli* cell membrane up to a critical threshold level. Site-directed mutagenesis of the holin gene was used to generate phage variants that vary in their timing of lysis from 30 to 190 min. Observation of the lysis times of single cells reveals an intriguing finding – the noise in lysis timing first decreases with increasing lysis time to reach a minimum, and then sharply increases at longer longer lysis times. A mathematical model with stochastic expression of holin together with dilution from cell growth was sufficient to explain the non-monotonic noise profile, and identify holin accumulation thresholds that generate precision in lysis timing.

## INTRODUCTION

The inherent probabilistic nature of biochemical reactions and low copy numbers of molecules involved results in significant random fluctuations (noise) in protein levels inside isogenic cells inhabiting the same environment (1–6). While the origins of stochastic gene expression have been extensively studied across organisms, how the noisy expression of key regulatory proteins impacts the timing of intracellular events is not well understood (7).

To address this knowledge gap, we use the bacteriophage λ as a model system for studying event timing at the single cell level. Here, an easily observable event (cell lysis) is the result of the expression and accumulation of a single protein (holin) in the *Escherichia coli* cell inner membrane up to a threshold level (Figure 1A) (8,9). Once holin surpasses this critical threshold concentration, it nucleates to form large holes in the inner membrane, triggering events that result in the destruction of the cell and the release of phage progeny (8,9). Since holin nucleation and cell lysis are essentially simultaneous, holin can be said to be the timekeeper of the lysis event (10). Single-cell observations of holin-induced lysis allows calculation of both the mean and noise of lysis timing, where noise is quantified using a dimensionless metric, the coefficient of variation (standard deviation divided by the mean). Our prior work revealed incredible precision in lysis timing in the wildtype λ strain: lysis occurs on average at 65 min with a coefficient of variation of less than 5% (11,12).

**Figure 1.**
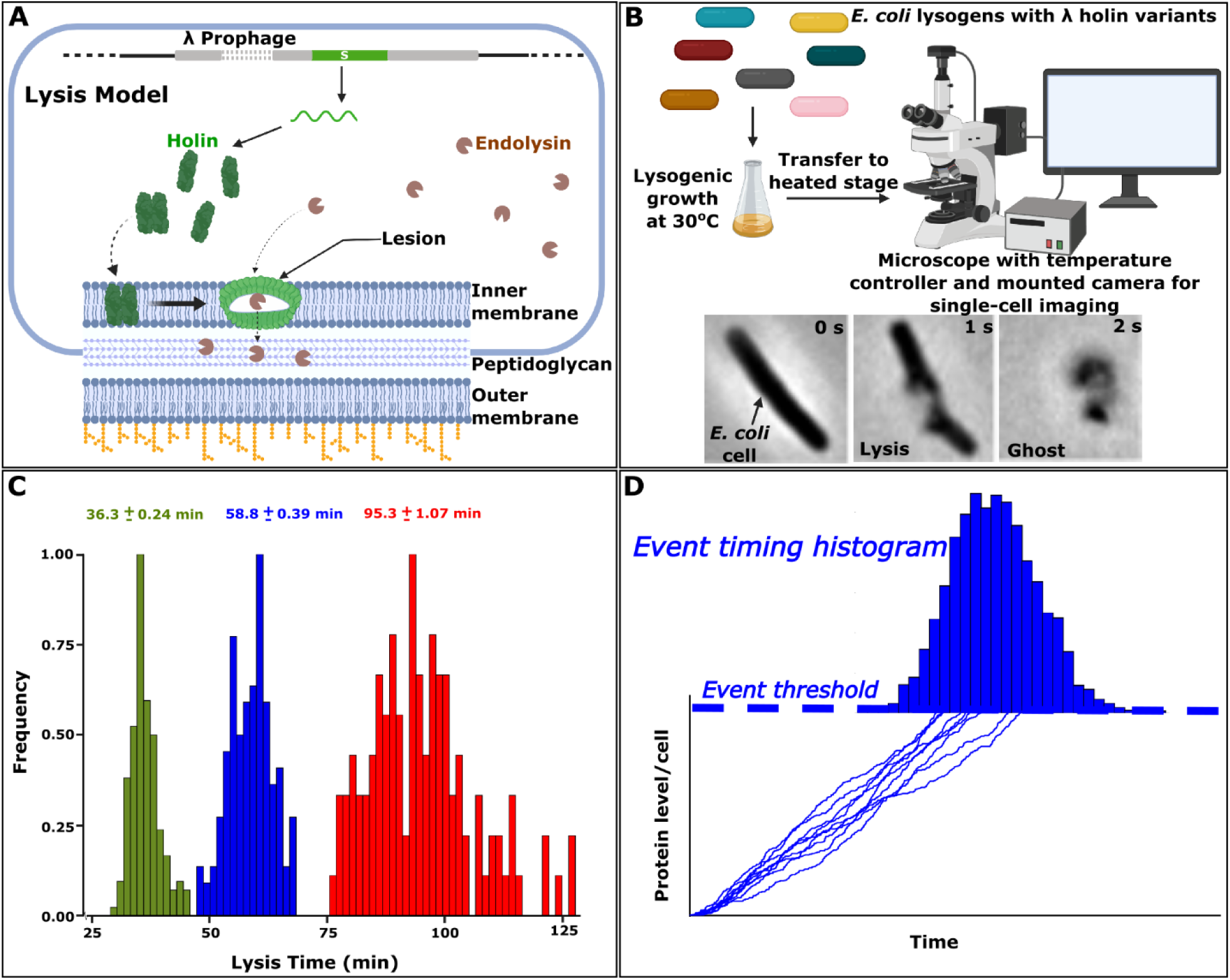
Phage lambda model system can be manipulated to study lysis time variation. **A)** Lysis model showing cell lysis mediated by λ holin. Accumulation of holin in the inner cell membrane results in lesions, which trigger cell lysis by allowing endolysin to access the *E. coli* cell wall. **B)** Lysogens were used for single-cell imaging and recording of lysis events. Inset panels track the lysis event occurring in a single cell after induction (original images courtesy of Ry Young). The third image shows the cell debris (ghost) after lysis. **C)** Lysis time distributions for three lysogens (∼100 cells) with different means. Mean ± SEM values are shown for each distribution. **D)** Cartoon showing stochastic accumulation of holin over time, and lysis time is the first-passage time (FPT) for holin levels to reach a critical threshold. Since expression is stochastic, the threshold is reached at different times in different cells.

Bacteriophage λ’s lysis timing provides an elegant experimental model system to probe the impact of parameter perturbations (e.g., of transcription rate, translation rate, etc.) on precision in the timing of intracellular events. The key objective here is to investigate how manipulation of the lysis timing threshold affects the noise in lysis timing. To this end, we systematically altered the amino acid sequence of the holin protein in order to shift (both increase or decrease) the lysis timing threshold (11,13). These amino acid sequence changes may affect holin structure, dimerization and/or oligomerization potential, and/or membrane insertion capacity. Sequence alterations that inhibit holin’s ability to pass into the inner membrane may, for example, increase the threshold, whereas alterations increasing holin-holin affinity may decrease the threshold. This contribution studies the effects of these alterations on noise in lysis timing both experimentally and via mechanistic mathematical models to uncover an intriguing insight – precision in timing is enhanced at an intermediate threshold.

## RESULTS and DISCUSSION

Timing of intracellular events is often studied using a first-passage time (FPT) framework that captures the first time a random process crosses a threshold (14–16). In our prior work, we formulated lysis timing as an FPT problem (11,13). Here, the start of transcription from λ’s late promoter results in stochastic accumulation of holin within the host cell, and lysis is triggered when the total cellular holin concentration reaches a critical threshold (Fig. 1D). Our mathematical analysis predicted that noise in timing is inversely proportional to the threshold (11,13). The logical progression of this work is to verify this prediction through experimental manipulation of the lysis threshold by altering the holin sequence.

It is important to point out that the holin gene *S* of wildtype λ has two translation initiation sites. Gene expression results in the production of two proteins, holin and antiholin, in a 2:1 ratio ensuring excess holin (17). Antiholin has two extra residues, a methionine and a leucine, and acts antagonistically to holin, which has the lysis function (18,19). To remove any confounding effects of antihloin on the lysis timing, we introduced mutations into a strain of λ where antiholin expression has been abolished via the M1L mutation (Table 1).

**Table 1.** Mean and noise in first-passage time (FPT) of isogenic *E. coli* λ lysogens. Single cell FPTs were calculated by subtracting 15 min from the recorded lysis times to account for the time delay between induction and start of transcription from the λ’s late promoter.

| Strain | Holin Mutations<sup>a</sup> | n<sup>b</sup> | Mean FPT (min)<br>(95% confidence interval)<sup>c</sup> | FPT CV<sup>2</sup><br>(95% confidence interval)<sup>c</sup> |
| --- | --- | --- | --- | --- |
| JJD3 (WT) | None | 120 | 48.83<br>(48.14–49.54) | 0.007<br>(0.0051–0.0089) |
| JJD5 (S105) | M1L | 140 | 43.77<br>(43.02–44.53) | 0.011<br>(0.0089–0.0137) |
| JJD246 | M1L/H7D | 114 | 54.02<br>(52.86–55.16) | 0.013<br>(0.0097–0.0179) |
| JJD248 | M1L/F94C | 121 | 41.01<br>(40.23–41.8) | 0.012<br>(0.0094–0.0157) |
| JJD251 | M1L/A99V | 128 | 25.89<br>(25.24–26.52) | 0.02<br>(0.0158–0.0237) |
| JJD253 | M1L/L10M | 149 | 26.19<br>(25.51–26.86) | 0.026<br>(0.0183–0.0337) |
| JJD388 | M1L/L25V/<br>N37H | 99 | 175.09<br>(163.8–186.1) | 0.10<br>(0.0627–0.1492) |
| JJD390 | M1L/A11G/<br>Y31H | 143 | 170.65<br>(159.9–181.3) | 0.13<br>(0.1116–0.1581) |
| JJD391 | M1L/A16G/<br>K92Q | 115 | 125.94<br>(122–130.1) | 0.031<br>(0.0156–0.0488) |
| JJD404 | M1L/I21V | 116 | 15.18<br>(14.63–15.74) | 0.038<br>(0.0253–0.052) |
| JJD405 | M1L/V45G | 118 | 17.13<br>(16.7–17.56) | 0.02<br>(0.0147–0.0259) |
| JJD411 | M1L/D85G | 91 | 161.76<br>(152.4–171.4) | 0.086<br>(0.0699–0.1042) |
| JJD413 | M1L/I87L | 158 | 21.32<br>(20.86–21.78) | 0.020<br>(0.0157–0.025) |
| JJD414 | M1L/L90I | 174 | 29.96<br>(29.2–30.7) | 0.029<br>(0.0229–0.0348) |
| JJD415 | M1L/I91T | 166 | 29.11<br>(28.37–29.85) | 0.03<br>(0.0241–0.0358) |
| JJD426 | M1L/G38S | 100 | 140.26<br>(134–146.5) | 0.053<br>(0.0374–0.0695) |
| JJD428 | M1L/G39S | 111 | 80.31<br>(78.23–82.38) | 0.02<br>(0.015–0.0252) |
| JJD432 | M1L/S89W | 132 | 92.74<br>(90.95–94.61) | 0.013<br>(0.0093–0.0176) |
| JJD434 | M1L/D8G | 104 | 76.09<br>(74.3–77.93) | 0.015<br>(0.01–0.025) |
| JJD436 | M1L/K92N | 138 | 85.99<br>(83.86–88.12) | 0.021<br>(0.0116–0.03) |

<sup>a</sup>Amino acid substitutions; <sup>b</sup>Number of cells observed; <sup>c</sup>95% CIs after bootstrapping (1,000 replicates); FPT, first-passage time; WT, wildtype.

Site-directed mutagenesis was used to introduce one or two nucleotide substitutions into plasmids bearing the *S105* holin allele. The resulting plasmids were used to generate a library of *E. coli* lysogens differing from the *S105* mutant by one or two amino acid substitutions in the holin gene. The optical densities of thermally induced batch cultures of these lysogens were tracked to determine their lysis times (unpublished data). For this study, we selected a subset of twenty holin mutants spanning a wide range of mean lysis times (Table 1). For each mutant strain, we thermally induced and recorded single cell lysis events for ∼100 cells using a microscope-mounted, temperature-controlled perfusion chamber (Figure 1B and C, movie file Lysis.asf). The mean lysis times calculated using both the batch culture and single-cell recordings were strongly correlated (Figure S1).

Using a subset of lysogens, we verified that the holin mutations had no effect on holin expression via Western blot assays of holin levels in whole-cell extracts (Figure S2). Therefore, any effects on lysis timing can be attributed to shifting of lysis threshold as a result of the amino acid changes in the holin gene. These changes in holin may affect lysis timing by altering holin-holin affinity, holin accession to the inner membrane, and/or holin nucleation within the inner membrane. To investigate these possibilities further, we compared levels of different holin mutants in the cell membrane. Interestingly, a lysogen with short lysis time showed almost five-fold higher mutant holin levels in the membrane compared to wildtype holin (Figure S2). Contrarily, a lysogen with long lysis time showed mutant holin levels comparable to wildtype holin. In the latter case, the mutant holin might be impaired in the formation of membrane lesions required for lysis, and thus delaying lysis. These results suggest that the quality of holin may directly affect the quantity of holin in the membrane and/or its ability to form the membrane lesions critical for lysis. Further biochemical studies may reveal how structural changes in holin affect the different steps leading to cell lysis.

Next, we quantified single-cell FPTs by subtracting 15 min from the recorded lysis times. This 15 min duration accounts for the time delay between lysogen induction and start of transcription from λ’s late promoter (20,21). Recall that simple models predict the noise in FPT to monotonically decrease with increasing lysis threshold (and hence, increasing mean FPT). Computations of both the mean and noise in FPTs across holin mutants as illustrated in Table 1 reveals an intriguing result – for short-lysis strains decreasing the lysis threshold increases the noise consistent with our previous model. By contrast, the data for long-lysis strains *contradicts* our simple model; increasing the lysis threshold *increases* the FPT noise level (Fig. 2). The concave-up shape of the plot implies that noise in FPT is minimized at an intermediate threshold. Interestingly, the wildtype λ genotype resides near the base of this plot suggestion that buffering noise in lysis timing is ecologically relevant, and is consistent with the existence of optima in lysis timing (12,22–24).

**Figure 2.**
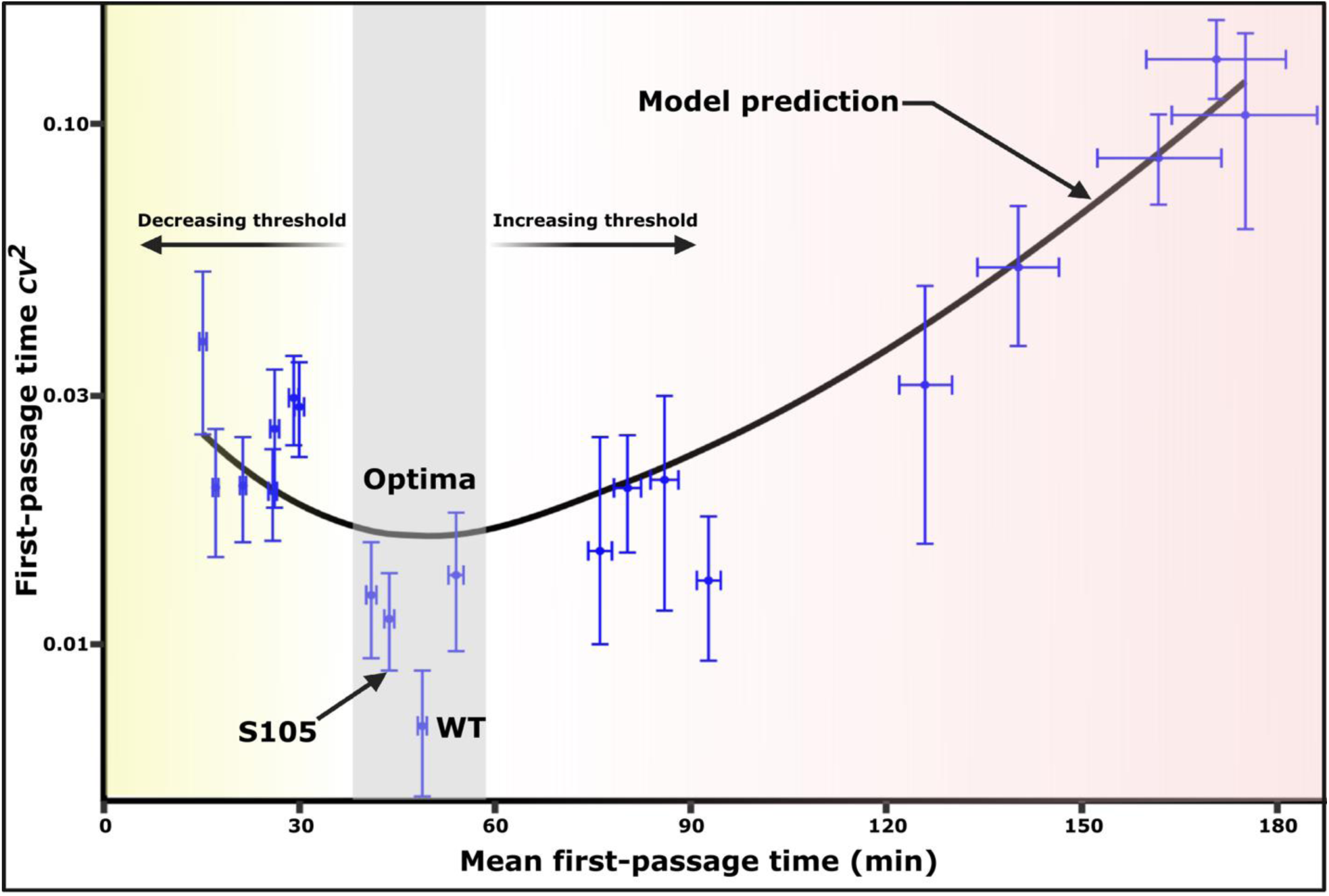
Noise in lysis timing is minimized at intermediate threshold. Noise in first-passage time (FPT) as quantified using the coefficient of variation squared (*CV*^*2*^) is shown plotted against mean FPT across holin mutants. Each point represents an isogenic λ strain with amino acid substitutions (Table 1) affecting the lysis timing threshold. These mutants show changes in FPT and *CV*^*2*^ consistent with the model prediction (black line, eq. 2). Eq. 2 was fitting to the data considering a 40 min cell doubling time (i.e., *E. coli* growth at 30°C), with a single-fitting parameter *CV*_*x*_, which was estimated to be *CV*_*x*_ = 0.05. Threshold is optimal at the base of the plot where the noise is minimized. WT, λ strain with wildtype *S* gene; S105, λ strain bearing the *S105* allele (holin); error bars, 95% CIs after bootstrapping (1,000 replicates).

To explain this non-monotonic noise profile, we developed an expanded model for noisy holin expression that explicitly considers dilution from cellular growth (see section S3 in SI). More specifically, as has been shown for *E. coli* genes (4,25,26), we consider holin expression occurring in stochastic bursts with holin dilution occurring between two successive burst events. Subsequent analysis of the model predicts the mean FPT as (details in SI)

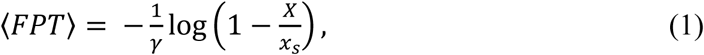

where *γ* is the cellular growth rate, *X* is the lysis threshold, and *x*_*s*_ is the steady-state mean holin concentration reached after a long time if there was no lysis. Moreover, the noise in FPT was derived as

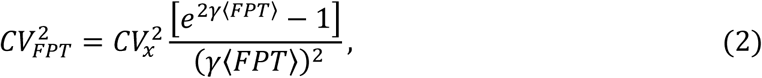

where the constant 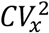 quantifies the extent of stochasticity in holin expression. Intriguingly, this formula predicts the timing noise to vary non-monotonically with the mean FPT and provides an excellent fit to the data (black line in Fig. 2). A key insight from (2) is that the noise is minimal when the threshold is 55% of the steady-state holin concentration *x*_*s*_.

In summary, our study uncovers mechanisms for generating precision in the timing of cellular events given the unavoidable constraints of stochastic gene expression and dilution from cellular growth. We show that genetic variation in event-timing noise exists, and hence is potentially a target of selection. This prediction can be tested in future work by comparing the fitness of mutant strains with same mean lysis timing but different noise levels.

## MATERIALS AND METHODS

### Bacterial and phage strains

All the bacteria and plasmids used in this study are listed in Table 2. *E. coli* lysogens were cultured in lysogeny broth (LB) at 30°C with rotary shaking (220 rpm).

**Table 2.**
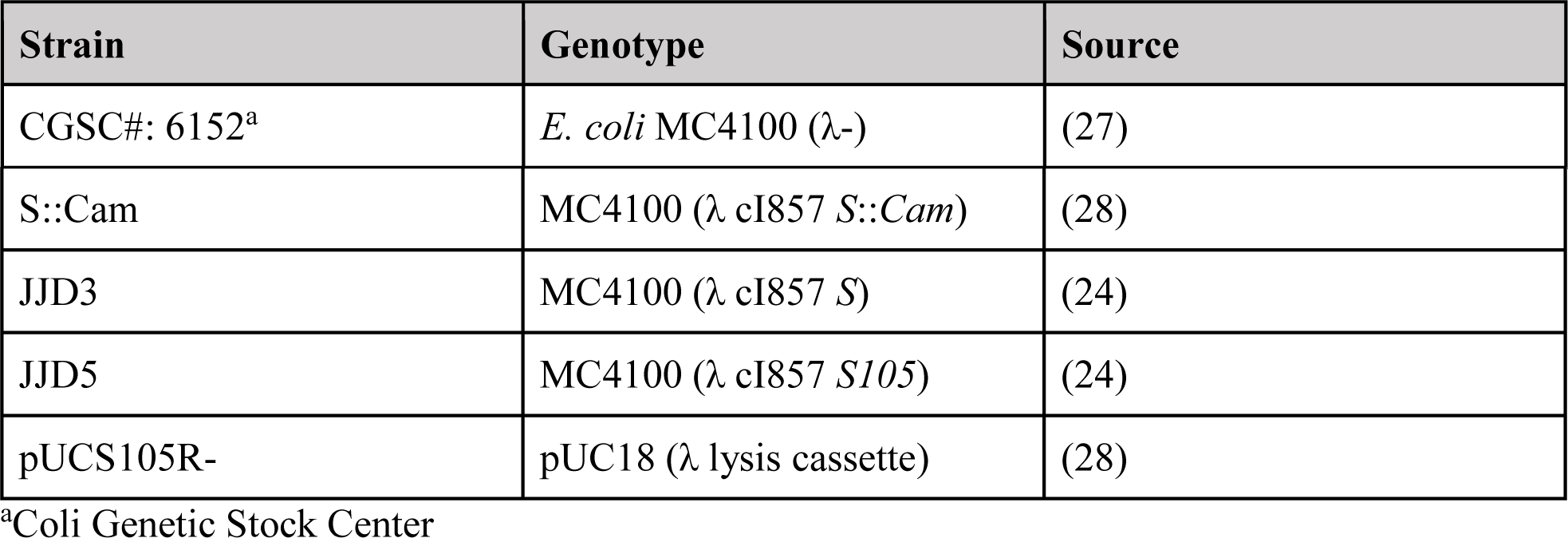
List of bacterial strains and plasmids used in this study.

| Strain | Genotype | Source |
| --- | --- | --- |
| CGSC#: 6152<sup>a</sup> | *E. coli* MC4100 ($\lambda^-$) | (27) |
| S::Cam | MC4100 ($\lambda$ cI857 S::Cam) | (28) |
| JJD3 | MC4100 ($\lambda$ cI857 S) | (24) |
| JJD5 | MC4100 ($\lambda$ cI857 S105) | (24) |
| pUCS105R- | pUC18 ($\lambda$ lysis cassette) | (28) |

<sup>a</sup>Coli Genetic Stock Center

### Construction of plasmids with mutations in holin

Briefly, site-directed mutagenesis was used to generate a panel of mutant λ phages with one or two base pair substitutions in the *S105* allele of the holin gene. Plasmid pUCS105R-carries the λ lysis cassette with the *S105* allele, which has a Leu (CUG) codon in place of the Met1 codon of the *S* gene. This plasmid was used as a template for PCR (Pfu DNA polymerase; Promega, Madison, WI) using megaprimers consisting of 30 to 45-nucleotide homology flanking the altered nucleotides. After *DpnI* treatment to digest the original template, the resulting plasmids were transformed into MC4100 (λ cI857 *S*::*Cam*) cells. The cells were spread on LB + Amp (100 μg ml^-1^) plates and incubated at 30°C until colonies were visible. Some of the plasmids thus constructed were further used as templates to generate double mutants.

### Transferring mutant holin from the plasmid into the λ phage genome

Transformed MC4100 (λ cI857 *S*::*Cam*) cells were grown in 3 ml LB supplemented with ampicillin (100 μg ml^-1^) at 30°C in a rotary shaker. For thermal induction of prophages, the cultures were transferred to a shaker at 42°C for 15 min and then 37°C until lysis. The resulting lysate was plated with MC4100 cells to obtain plaque-forming phages resulting from recombination between the prophage and the plasmid. To obtain lysogens, phages obtained from the plaques were used to infect 100 μl of saturated MC4100 culture for 30 min. To this culture, 1 ml LB broth was added and further incubated at 30°C in a rotary shaker for 1 h. A 100 μl aliquot of this mixture was spread on LB plates supplemented with kanamycin (50 μg ml^-1^), and incubated overnight at 30°C. The lysogens were further screened for ampicillin resistance followed by DNA sequencing to confirm the nucleotide substitutions.

### Single-cell lysis time determination

The protocol for determining single-cell lysis times has been described previously (6). Briefly, lysogens were grown overnight in LB at the permissive temperature of 30°C. Overnight cultures were diluted 100-fold and grown to A550 = 0.3–0.4 in a 30 °C shaking incubator. A 200-µl aliquot of the exponentially growing culture was chemically fixed to a 22 mm square glass coverslip, which was pretreated with 0.01% poly-L-lysine (mol. wt. 150 K–300 K; Millipore Sigma, St. Louis, MO) at room temperature for 10 min, and applied to a perfusion chamber (RC-21B, Warner Instruments, New Haven, CT). After assembly, the perfusion chamber was immediately placed on a heated platform (PM2; Warner Instruments, New Haven, CT), which was mounted on an inverted microscope stage (TS100, Nikon, Melville, NY), and infused with heated LB at 30°C (Inline heater: SH-27B, dual channel heating controller: TC-344B; Warner Instruments, New Haven, CT). The chamber temperature was spiked to 42°C for 20 min, then maintained at 37°C until ∼95% lysis was observed. Videos were recorded using an eyepiece camera (10X MiniVID™; LW Scientific, Norcross, GA), and the lysis times of individual cells were visually ascertained using VLC™ media player. Lysis time was defined as the time required for a cell to disappear after the temperature was increased to 42 °C.

## ACKNOWLEDGEMENTS

We are grateful for intellectual input from Khem Gusingha, Cesar Vargas-Garcia, and past and present members of the Dennehy Lab. We thank Ryland Young (Texas A&M) for stimulating conversations about phage λ, and for some of the phage and bacterial strains described herein.

