## Supplementary Information for "Optimum Threshold Minimizes Noise in Timing of Intracellular Events"

### S1. Lysis time determination of batch cultures using a plate reader

After sequence confirmation, the lysogens were first heat-induced in batch cultures to assess their lysis times. A 5-µL aliquot of overnight cultures was mixed with 1 mL LB in 24-well plates. Following growth at 30°C for 2 h, the plates were shifted to a 42°C water bath (time 0 for lysis time) for 15 min. After heat induction, the plates were shifted to a pre-warmed plate reader (Synergy™ HT, BioTek® Instruments, Inc., Vermont, USA) at 37°C, which measures $A_{550}$ of the culture every 2 min. This protocol was repeated in triplicate for all lysogens. The complementary cumulative distribution function of the normal distribution was used to fit the $A_{550}$ outputs generated by the plate reader. The estimated mean and standard deviation were defined as the lysis time and spread respectively. FPTs estimated using both batch culture and single-cell recordings were strongly correlated (Figure S1).

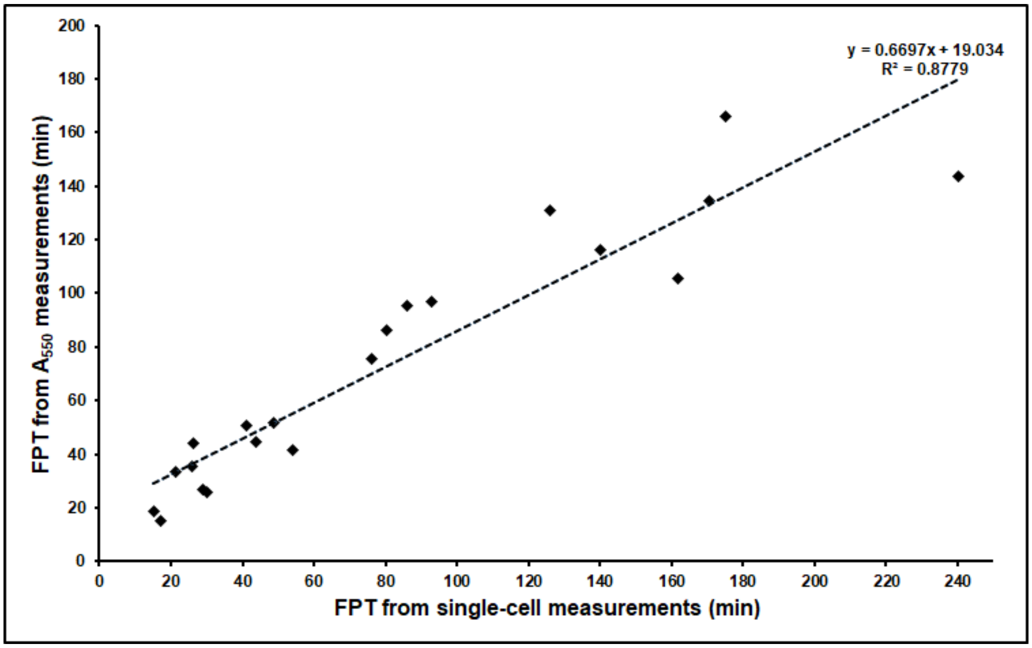


**Figure S2.** The FPT measurements using $A_{550}$ and single-cell recordings were strongly correlated.

### S2. Holin expression

We extracted holin from whole cells or from cell membranes to compare the relative holin levels in different mutant strains. An exponentially growing culture ($A_{600}$ ~0.4) at 30°C was induced at 42°C for 20 min. A 5 mL aliquot of the culture was then immediately centrifuged to pellet the cells. The pellets were mixed with 2× SDS-PAGE sample buffer, heated at 100°C for 5 min, and loaded on a 4-20% TruPAGE™ precast gel (Sigma-Aldrich, St. Louis, MO, USA). Another 4 mL aliquot of the culture was heat induced for 30 min and then sonicated to disrupt the membranes. The membranes were collected by centrifugation at 100,000 × g for 1 h. The pelleted membranes were mixed with 40 µl of ME buffer (10 mM Tris Cl [pH 8.0], 35 mM $MgCl_2$, 1% Triton X-100) by shaking on a platform shaker for 2 h at 25°C. The extracted samples were centrifuged at 100,000 × g for 30 min to pellet the insoluble fraction. The membrane extracts were mixed with 2× SDS-PAGE sample buffer, heated at 100°C for 3 min, and loaded on to precast gels. After electrophoresis, Western blotting was used to detect holin using a primary antibody (1:1000) raised in rabbits. A secondary antibody (donkey anti-rabbit polyclonal antibody conjugated to horseradish peroxidase [SA1-200, ThermoFisher Scientific, Waltham, MA, USA]) was used at a dilution of 1:1000 dilution, and the blot was developed as per the manufacturer's directions. An average of three preparations was used to estimate holin levels.

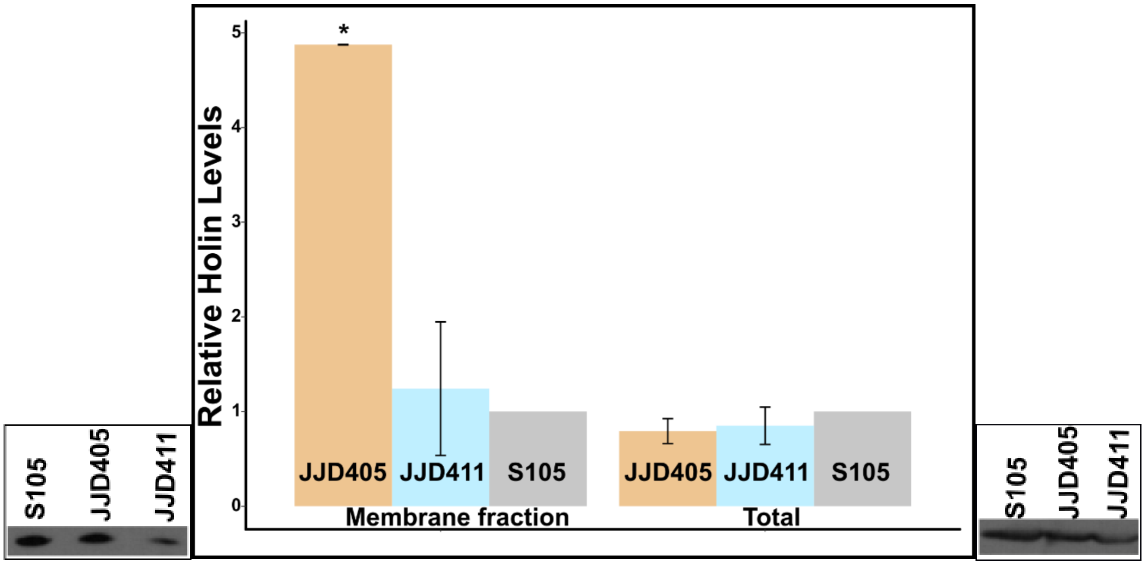


**Figure S2.** Total holin levels from whole-cell extracts and membrane fractions. The left and right panels show Western blots of membrane fractions and whole-cell extracts respectively. Bands represent holin extracted from strains S105 (mean LT = 58 min), JJD405 (32 min), and JJD411 (176 min). * = $p < 0.05$, t-test; error bars are SEM.

### S3. Calculation of noise in the first passage time

We modeled the expression of holin occurring in intermittent bursts, with burst events arriving as a Poisson process with rate $k$. Whenever a burst occurs, the total cellular concentration of holin $x(t)$ at time $t$ increases by a random amount $b$:

$$x(t) \mapsto x(t) + b, \qquad S1$$

where $b$ is drawn from an arbitrary positively-valued probability distribution with the first and second moments $\langle b \rangle$ and $\langle b^2 \rangle$, respectively. The first moment $\langle b \rangle$ represents the mean burst size per unit volume. Between two consecutive bursts, the concentration diluters from cell growth as per the following the deterministic dynamics:

$$\dot{x}(t) = -\gamma x(t) \qquad S2$$

where $\gamma$ is the cellular growth rate. For this hybrid system with stochastic bursts interspersed by first-order decay, the time evolution of the first and second-order moments of $x(t)$ are given by

$$\frac{d\langle x\rangle}{dt} = k\langle b\rangle - \gamma\langle x\rangle, \qquad S3$$

$$\frac{d\langle x^2\rangle}{dt} = k\langle b^2\rangle + 2k\langle b\rangle\langle x\rangle - 2\gamma\langle x^2\rangle, \qquad S4$$

(1,2,3). Solving the above differential equations, we get the mean $\langle x\rangle$ and variance $\langle x^2\rangle - \langle x\rangle^2$ of the holin concentration as a function of time $t$, assuming there is no holin at the onset of the protein synthesis;

$$\langle x\rangle = \frac{[1 - e^{-\gamma t}]\, k\,\langle b\rangle}{\gamma} \qquad S5$$

$$\langle x^2\rangle - \langle x\rangle^2 = \frac{[1 - e^{-2\gamma t}]\, k\,\langle b^2\rangle}{2\,\gamma}. \qquad S6$$

We formulate the lysis time as the first-passage time

$$FPT = \min\{t\colon x(t) \geq X | x(0) = 0\}, \qquad S7$$

or the first time the holin concentration reaches a critical threshold level $X$, and quantify the noise in the first-passage time using the coefficient of variation squared,

$$CV_{FPT}^2 = (\langle FPT^2\rangle - \langle FPT\rangle^2)/\langle FPT\rangle^2,$$

where $\langle FPT\rangle$ and $\langle FPT^2\rangle$ are the first two moments of $FPT$. $CV_{FPT}^2$ is related to the fluctuations in the holin concentration as per

$$CV_{FPT}^2 \approx \frac{\langle x^2\rangle - \langle x\rangle^2}{\langle FPT\rangle^2}\left(\frac{d\langle x\rangle}{dt}\right)^{-2}\Bigg|_{t=\langle FPT\rangle}, \qquad S8$$

(4). From eq. S5 the mean first-passage time is obtained as

$$\langle FPT\rangle = -\frac{1}{\gamma}\log(1-\alpha), \text{ with } \alpha = \frac{X}{x_s}. \qquad S9$$

Here $x_s$ denotes the steady-state mean holin concentration and is given by,

$$x_s = \langle x(t\to\infty)\rangle = \frac{k\,\langle b\rangle}{\gamma},$$

with the underlying assumption in eq. S9 being that the threshold for lysis $X$ is less than $x_s$. Using equations (S5), (S6), and (S8), we write down the formula for the noise in $FPT$,

$$CV_{FPT}^2 = CV_x^2\frac{\alpha\,(2-\alpha)}{[(1-\alpha)\ln(1-\alpha)]^2}, \qquad S10$$

where, $CV_x^2$ is the coefficient of variation squared for the holin concentration at steady state

$$CV_x^2 = \lim_{t \to \propto} \frac{\langle x^2 \rangle - \langle x \rangle^2}{\langle x \rangle^2} = \frac{\langle b^2 \rangle}{2\langle b \rangle x_s}$$

and quantifies the extent of stochasticity in holin expression. The above formula can be rewritten in terms of $\langle FPT \rangle$ as,

$$CV_{FPT}^2 = CV_x^2 \frac{\left[e^{2\gamma\langle FPT \rangle} - 1\right]}{(\gamma\langle FPT \rangle)^2} \qquad S11$$

and varies nonmonotonically with the mean FPT consistent with experimental data. The optimal value of the mean FPT (in the unit of $\gamma^{-1}$), where noise is minimum is $\gamma\langle FPT \rangle \approx 0.8$. The corresponding value of the threshold (in the unit of steady state concentration) is $\alpha = \frac{X}{x_s} \approx 0.55$.
